# Mechanistic characterization of the antiviral effects of Nordihydroguaiaretic Acid against West Nile Virus

**DOI:** 10.1101/2025.05.10.653272

**Authors:** Florencia Martinez, Lucia María Ghietto, Giuliana Lingua, Pedro Ignacio Gil, Juan Javier Aguilar, Tomas Isaac Gomez, Juliana Marioni, Susana Carolina Núñez-Montoya, Brenda Salome Konigheim

**Affiliations:** Universidad Nacional de Córdoba, Facultad de Ciencias Médicas, Instituto de Virología “Dr. J. M. Vanella”, Cdad. Universitaria, X5000HUA Córdoba, Argentina; Consejo Nacional de Investigaciones Científicas y Técnicas (CONICET); Universidad Nacional de Córdoba, Facultad de Ciencias Químicas, Departamento de Ciencias Farmacéuticas. Haya de la Torre y Medina Allende, Ciudad Universitaria. X5000HUA Córdoba, Argentina; CONICET - Unidad de Investigación y Desarrollo en Tecnología Farmacéutica (UNITEFA). Haya de la Torre y Medina Allende, Ciudad Universitaria. X5000HUA Córdoba, Argentina

**Keywords:** Arboviruses, natural product, lignan, antiviral activity, cytoskeleton

## Abstract

Mosquito-borne Arboviruses, including Alphaviruses and Flaviviruses, are responsible for millions of human infections worldwide. The absence of specific antivirals for arbovirus infections underscores the urgent need for novel research to develop effective treatments. Natural products have emerged as promising candidates. Nordihydroguaiaretic acid (NDGA), a natural lignan predominantly found in Larrea spp., has shown antiviral activity against several viruses, including Dengue. This study aimed to investigate the antiviral potential of NDGA against West Nile virus (WNV), assessing its mechanism of action and cellular interactions. Using plaque-forming unit reduction and immunofluorescence assays in LLC-MK2 cells, we determine the reduction in viability of WNV viral particles at different times post-infection. The potential involvement of the sterol regulatory element-binding proteins (SREBP) and 5-lipoxygenase pathways in WNV infection was evaluated. Additionally, we analyzed the cytoskeleton integrity in treated cells. NDGA, had the greatest inhibitory effect on WNV replication when added 1 and 2 h post-viral internalization (PI), losing activity when added after 3 h PI, thus suggesting it targets early viral infection events. The SREBP pathway seems to be involved in the inhibition of WNV by NDGA, likely due to its lipogenic activity. Furthermore, the antiviral activity of NDGA could be related to its ability to modify the cytoskeleton of LLC-MK2 cells. These findings highlight NDGA potential as a therapeutic candidate against WNV and other flaviviruses. Plant metabolites are among the leading therapeutic resources globally. Further research is necessary to comprehensively assess the therapeutic promise of NDGA.

**Importance:** Mosquito-borne arboviruses pose significant global health threats due to their widespread distribution and the absence of specific antiviral treatments. The emergence of virulent strains and recurrent outbreaks underscore the urgent need for novel antiviral agents. Nordihydroguaiaretic acid, a natural phenolic compound, has shown promising antiviral activity against flaviviruses such as West Nile virus. Understanding nordihydroguaiaretic acid’s mechanism of action not only reveals potential therapeutic strategies but also highlights the broader importance of plant-derived compounds in combating viral infections. Our research focuses on further elucidating nordihydroguaiaretic acid’s antiviral effects against West Nile virus, aiming to contribute to the development of effective therapies. Continued investigation of nordihydroguaiaretic acid’s therapeutic potential is essential for advancing antiviral drug discovery and addressing the persistent challenges posed by mosquito-borne diseases worldwide.

## Introduction

Arboviruses, arthropod-borne viruses, represent a substantial threat to global health, manifesting in a range of diseases from mild febrile conditions to severe neurological complications. Since its emergence, West Nile virus (WNV) has proliferated throughout the Americas, leading to numerous outbreaks with significant economic repercussions (1). Such outbreaks have the potential to overwhelm healthcare systems, as evidenced by the SARS-CoV-2 pandemic (2). WNV, classified within the Flavivirus genus, is responsible for a seasonal arbovirus infection that can result in severe neurological disease. Its transmission cycle involves avian species as reservoirs, incidental hosts such as horses and humans, and the *Culex* spp. mosquitoes as the vector of infection (3). This ecological cycle facilitates the global dissemination of the virus. In addition to WNV, the Flavivirus genus encompasses other notable pathogens, including Dengue (DENV) and Zika virus, which share a similar structural architecture, indicating potential cross-reactivity with antiviral agents targeting these viruses.

In light of the pressing demand for effective antiviral therapies, there is currently no approved vaccine or specific pharmacological treatment available for WNV infections in humans (4). Medicinal plants and its metabolites have been highly valued for millennia as a rich source of therapeutic agents for managing viral disorders. Nordihydroguaiaretic acid (NDGA), a natural phenolic compound, sourced from *Larrea* spp., has been suggested for the treatment of various ailments, including cancer and chronic conditions such as diabetes, cardiovascular disease, and inflammatory disorders (5). Moreover, prior research has demonstrated the antiviral efficacy of NDGA against several arboviruses, indicating its potential as a broad-spectrum antiviral agent against arbovirus infections (6-9). Our findings reveal that NDGA exhibits both virucidal and antiviral effects against DENV1 *in vitro*, particularly within the initial 2 h following viral internalization. Through the application of qPCR and immunofluorescence techniques, it was confirmed that the reduction in infected cells would be associated with a decrease in viral replication, suggesting an intracellular mechanism of action. It is demonstrated that NDGA inhibits the endosome acidification, as its lysosomotropic effect impairs the viral uncoating process (8).

Elucidation of the molecular mechanisms underlying the antiviral activity of NDGA would provide new insights into virus-host interactions. The capacity of NDGA to modulate critical cellular pathways suggests its potential utility in the development of more effective therapeutics against arboviruses. In this work, we aim to continue further studying the effects of NDGA, specifically on WNV infection.

## Results

### WNV replication cycle

To characterize the different phases of the viral growth curve, the amount of infectious virus produced over time was evaluated by means of plaque assay in LLC-MK2 cells infected with WNV (MOI 1). Figure 1-A shows the duration of the different phases of the WNV growth curve. No increase in the infectious titer of the virus was detected in the extracellular medium during the initial 12 h PI (eclipse phase). Maximal viral yields were observed in the exponential or logarithmic phase, which lasted from 12 to 30 h PI. Finally, infected cell monolayers entered their stationary phase, displaying signs of lysis and cell death. The presence of viral proteins was first visualized by IF at 12 h PI in the perinuclear region, and after 24 h PI in a large area of the cytoplasm (Fig. 1-B).

**Figure 1.**
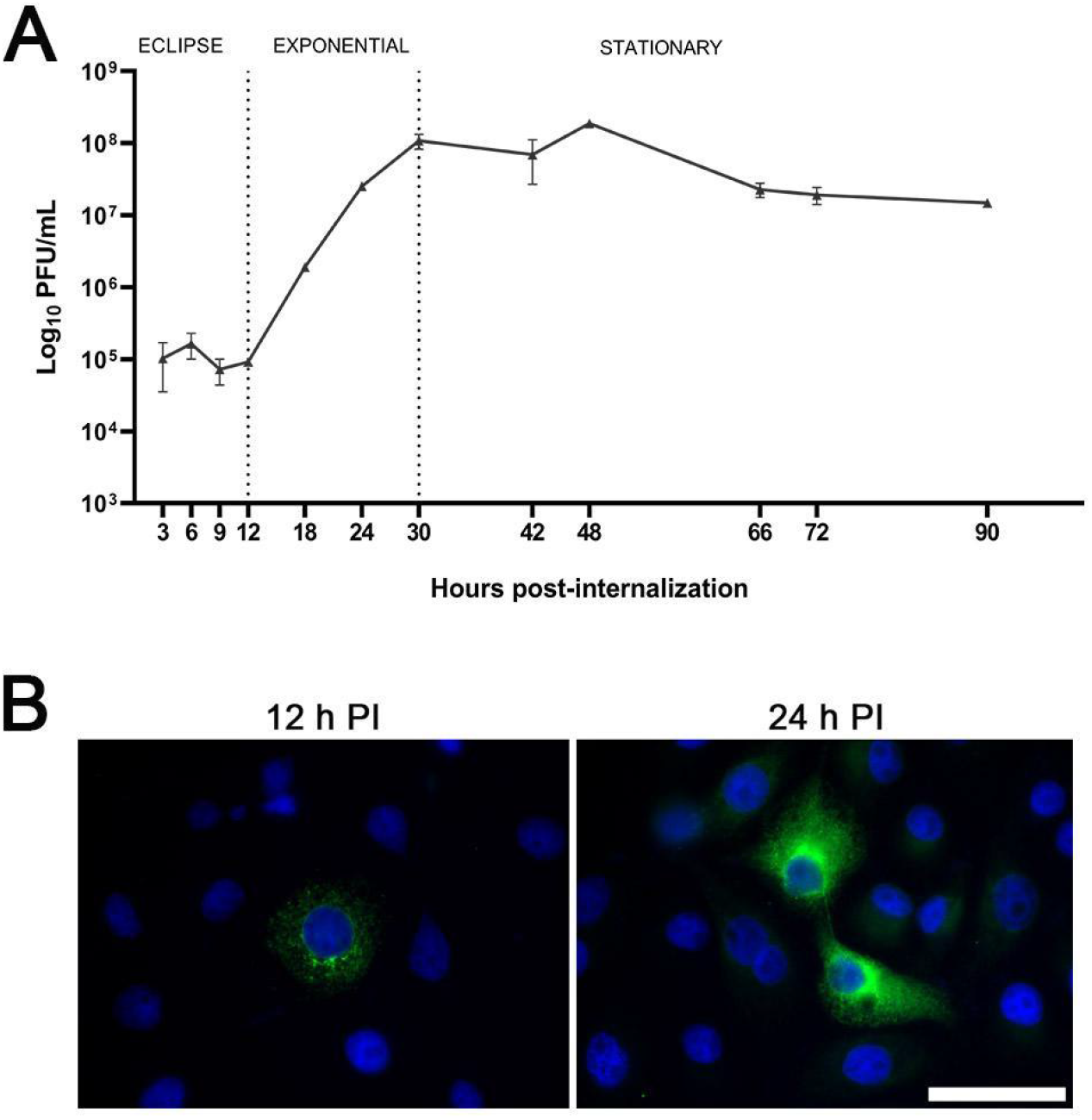
WNV replication cycle in LLC-MK2 cells. **A** WNV growth curve in LLC-MK2 cells. Monolayers were infected with MOI 1, and culture media were collected at different PI times (3, 6, 9, 12, 18, 24, 30, 42, 66, 72, and 90 h), and then analyzed for infectious viral particles using a standard plaque assay (log_10_ PFU/mL). Values shown viral titers (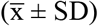) obtained by PFU assay in duplicate. **B** Representative images of WNV protein production in cultures at 12 and 24 h PI, evaluated by immunofluorescence assay. LLC-MK2 cells were fixed at 12 and 24 h PI and processed for IF microscopy using an anti-WNV primary antibody from serum and an anti-human FITC-IgG secondary antibody (green). Scale bar=20 μm.

### NDGA inhibits WNV during the first two hours postinternalization

To evaluate the antiviral activity of NDGA, the MNCC of 90.4 ± 5.9 μM for LLC-MK2 cells was established (9). NDGA was then tested at concentrations ranging from 2.5 to 90 μM. A dose-dependent inhibition of WNV replication was observed, with a maximum of 91.1% inhibition at 90 μM. The EC_50_ value, calculated from the dose-response curve (Fig. 2-A), was 29.8 ± 1.6 µM. The selectivity index (SI) was 3.9, indicating selective inhibition of WNV replication over LLC-MK2 cells cytotoxicity.

**Figure 2.**
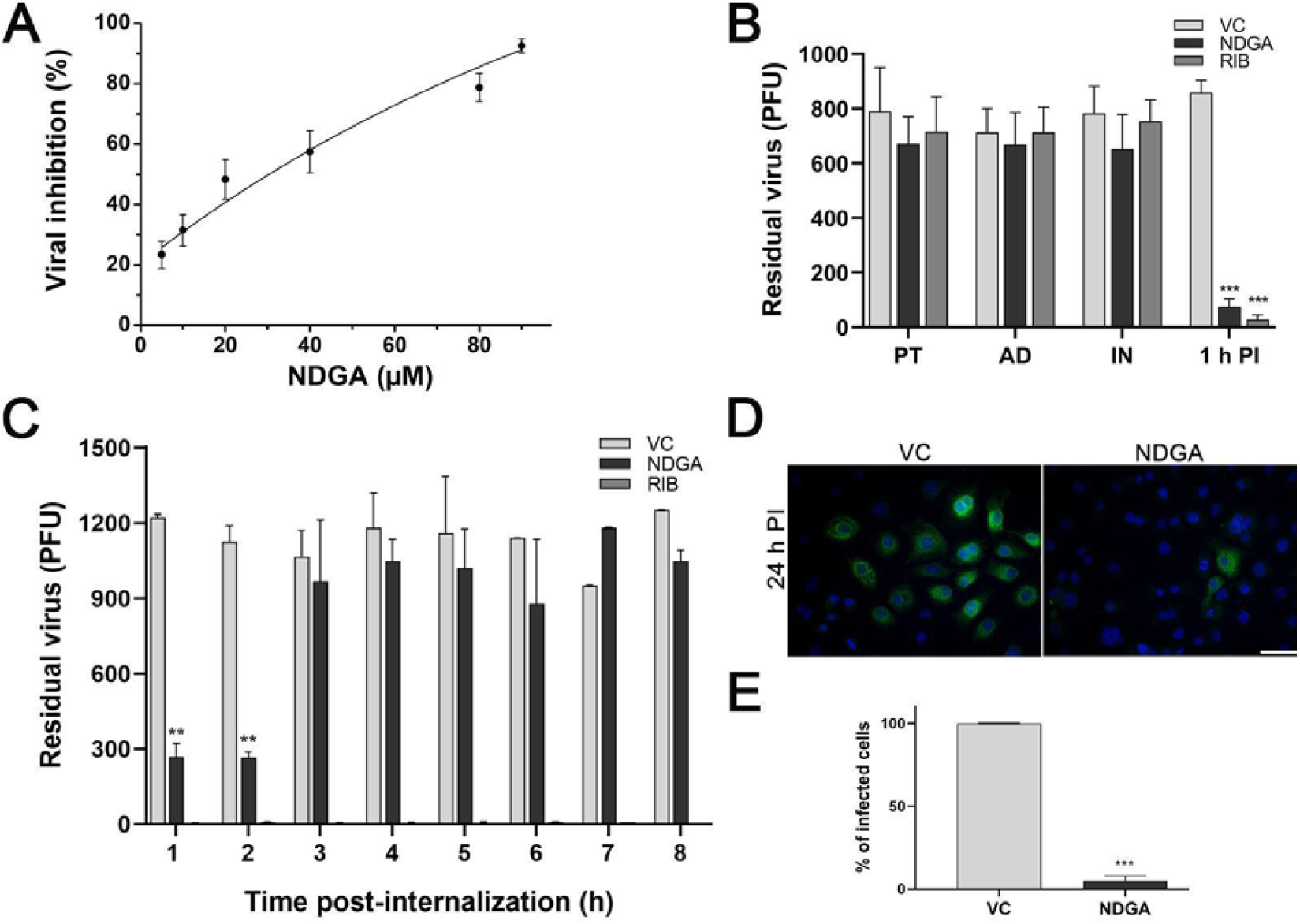
*In vitro* antiviral activity of NDGA against WNV. **A** Dose-response curve of NDGA antiviral activity against WNV. Results were fitted to a sigmoidal curve, R^2^ > 0.9. **B** Antiviral activity of NDGA and RIB at different infection times: pre-treatment (PT), adsorption (AD), internalization (IN), and 1 h PI. **C** Antiviral activity of NDGA and RIB at different PI times. **D** Representative images of cultures treated with NDGA or untreated and infected with a WNV (green) and at 1 h PI. Cell nuclei (blue). Scale bar: 20 μm. **E** Percentage of infected cells with WNV and treated with NDGA at 1 h PI against control (VC, 100%). Bars represent 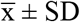 from three independent experiments performed in triplicate. Significance was calculated by one-way ANOVA test, followed by Tukey post-hoc analysis to allow comparison of specific group, with a *p* value ≤ 0.05 considered as statistically significant. \**p*< 0.05; \*\**p*< 0.01; \*\*\**p*< 0.001.

NDGA at CC_50_ concentration (115 μM) showed no significantly virucidal activity against WNV particles (2.3 x 10^9^ PFU/mL viruses treated with NDGA *vs*. 1.2 x 10^9^ PFU/mL in VC, *p*= 0.424), suggesting that the antiviral activity of this lignan is primarily due to its cellular effects.

To investigate the stage of the WNV life cycle targeted by NDGA, time-of-addition experiments were performed. NDGA treatment during pre-treatment, adsorption, and internalization did not significantly reduce WNV infection compared to the untreated control (Fig. 2-B). However, treatment initiated after viral internalization (1 h PI) resulted in inhibition of WNV replication (Fig. 2-B), suggesting that NDGA likely targets a post-viral internalization stage. To further define the window of NDGA antiviral activity, an additional experiment was conducted, NDGA was added at different PI times (1-8 h). NDGA treatment maintained a reduction in PFU compared to the untreated control up to 2 h PI,. This antiviral effect was no longer evident when NDGA was added at 3 h PI (Fig. 2-C).

The antiviral activity of NDGA against WNV was corroborated by immunofluorescence assays. Treatment with NDGA (90 μM) resulted in a significant reduction in the number of infected cells, as evidenced by a decrease in the expression of WNV proteins (Fig. 2-D). Quantification of infected cells revealed a statistically significant difference between the NDGA-treated group and the untreated group (*p*= 0.0005, Fig. 2-E), supporting the antiviral efficacy of NDGA.

#### NDGA’s antiviral activity: modulation of cellular factors as a potential mechanism

Antiviral drugs act through three main mechanisms: (a) directly targeting a viral protein, (b) inhibiting a cellular process essential for viral replication, or (c) modulating the host’s immune response (10). Given the broad-spectrum antiviral activity of NDGA against diverse viruses (6-9, 11-15) and its reported effects on cellular processes, such as antioxidant activity, inhibition of lipoxygenases (LOXs) and activation of signaling pathways (5), we hypothesized that this compound may act by modulating cellular factors (proviral factors), thus inhibiting WNV replication.

The antiviral effect of NDGA has been linked to the inhibition of lipogenesis (6, 7, 13). This compound is thought to exert its effect by downregulating the sterol regulatory element-binding protein (SREBP) pathway, which controls the expression of enzymes involved in sterol biosynthesis (13). Additionally, NDGA has been shown to inhibit the enzyme 5-LOX (5, 16), which is involved in the conversion of arachidonic acid to leukotrienes. Specific inhibitors of these pathways were employed to determine if they could replicate the antiviral effect of NDGA. Resveratrol, an inhibitor of SREBP1 the initial enzyme in the SREBP pathway (17), and caffeic acid, a selective inhibitor of 5-LOX (18), were chosen for this purpose. To ensure minimal cytotoxicity, concentrations of resveratrol and caffeic acid previously established as non-cytotoxic to LLC-MK2 cells were employed (9). Evaluation of their antiviral activity revealed that caffeic acid (200 μM) did not significantly affect WNV replication (Fig. 3-A). In contrast, resveratrol at 20 μM exhibited antiviral activity against WNV compared to untreated control, although this effect was weaker than that observed with NDGA at 90 μM (Fig. 3-A). These findings suggest that the antiviral activity of NDGA may be mediated, in part, by its inhibitory effects on the SREBP pathway.

**Figure 3.**
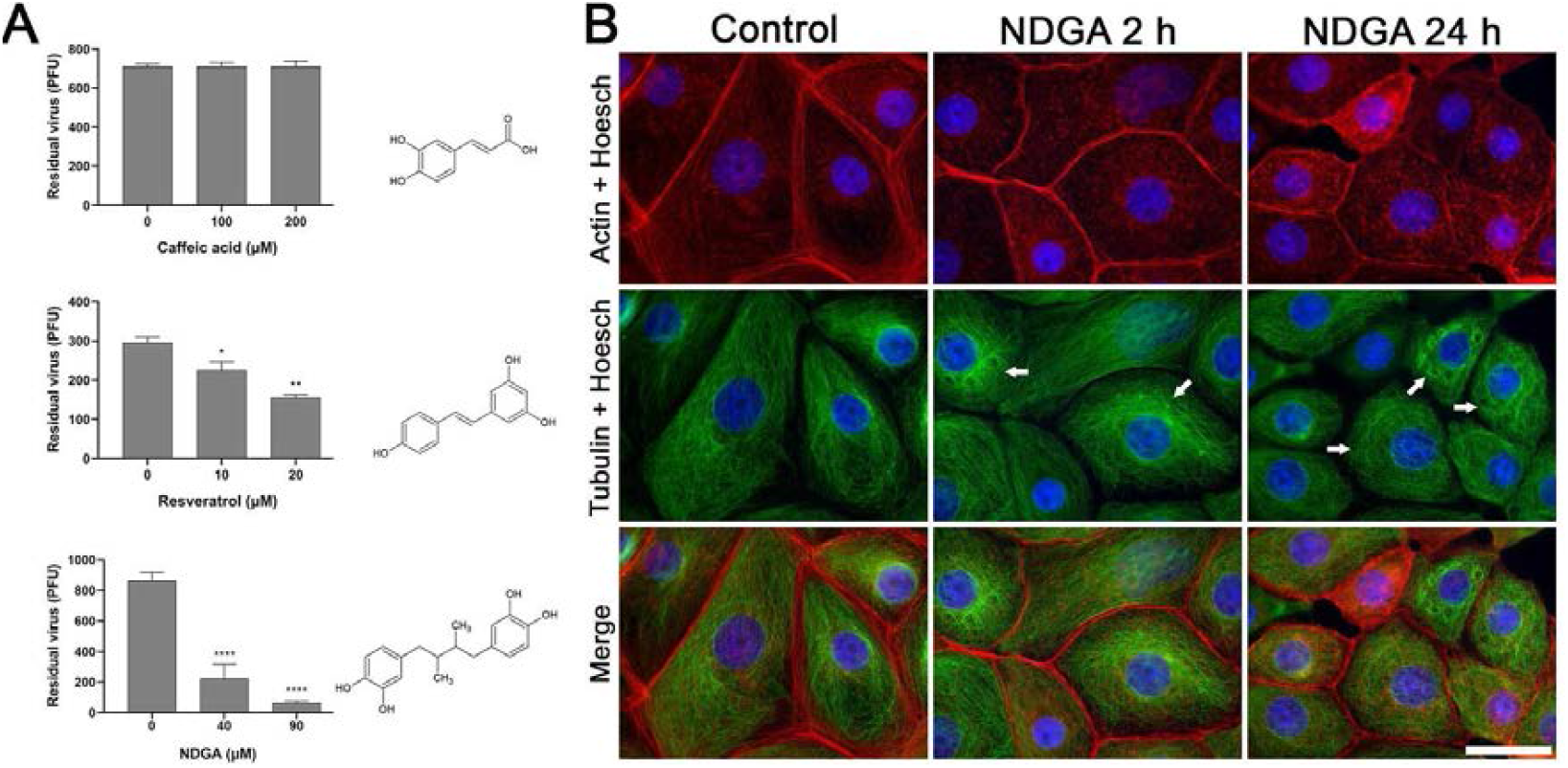
Characterization of the mechanism of action. **A** Chemical structure of inhibitors for the different cellular lipid metabolism pathways and antiviral activity of caffeic acid, resveratrol, and NDGA against WNV compared to the viral control (VC). Bars represent 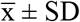 from three independent experiments performed in triplicate. Significance was calculated by one-way ANOVA test, followed by Tukey post-hoc analysis to allow comparison of specific group, with a *p* value ≤ 0.05 considered as statistically significant. \**p*< 0.05; \*\**p*< 0.01; \*\*\**p*< 0.001. **B** Time course changes in actin and tubulin cytoskeleton of LLC-MK2 cells. Cultures were treated with 90 μM of NDGA for 2 and 24 h and processed for IF. Actin and tubulin were visualized using phalloidin-rhodamine (red) and anti-α-tubulin antibody (green), respectively. Nuclei were stained with DAPI (blue). Note the progressive disorganization of the MT cytoskeleton in NDGA-treated cells (narrow). Scale bar: 20 µm.

In the viral infection cycle, the first hours of infection are crucial for viral uncoating (19). After internalization by clathrin-dependent endocytosis, trafficking of viral particles from early to late endosomes is dependent on cellular microtubules (MT)(20). We investigated whether NDGA treatment could alter the host cell cytoskeleton during the first 2 h post-treatment. This exploration aimed to elucidate a potential mechanism by which NDGA exerts its antiviral activity during this early stage of infection, focusing on its impact on the cellular cytoskeleton. For this, cell cultures were treated for 2 h with NDGA at 90 µM and processed for immunofluorescence. The consequence of NDGA treatment for 24 h was also evaluated to know the progress of the effect. We did not observe any changes in the actin cytoskeleton and, consequently, no alterations in cell morphology. However, NDGA treatment induced notable changes in MT organization (supplementary material). As shown in Figure 3-B, at 2 h post-treatment, the MT exhibited a loss of radial assembly from the microtubule-organizing center, redistributing into an irregular pattern towards the center of the cell. By 24 h treatment, we noted the appearance of circular rearrangements in the MT (narrow), a more pronounced effect. By 24 h of treatment, further changes were evident, characterized by the emergence of circular arrangements of MTs, indicating a progressive and distinctive effect induced by NDGA treatment. Morphological changes were quantified, and we observed that 82% and 65% of cells exhibited alterations at 2 h and 24 h post-incubation, respectively, compared to 25% in control cells (*p* < 0.0001).

## Discussion

Our research group has extensively explored the broad-spectrum antiviral potential of NDGA against different arboviruses, and identified it as a promising candidate for treating viral infections that currently lack approved antiviral therapies (8, 9, 14). This study confirmed the antiviral activity of NDGA against WNV and provided novel insights into its mechanism of action. We demonstrated that NDGA specifically targets cellular processes critical for viral replication (proviral factors), corroborating previous findings (7) and significantly expanding our understanding of this promising therapeutic candidate.

The growth kinetics of WNV in LLC-MK2 cells, assessed by plaque assay and immunofluorescence (IF) microscopy, revealed an eclipse phase of approximately 12 h, followed by a 30 h exponential growth phase. This growth curve, characterized by a distinctive eclipse phase and subsequent exponential viral release, is consistent with previously reported WNV growth kinetics in Vero B4 and C6/36 cells (21). The IF assay revealed the emergence of perinuclear viral protein localization at 12 h PI, which correlates with the onset of viral replication.

Consistent with our findings, previous studies have reported low *in vitro* cytotoxicity of NDGA in mammalian cells, with maximum non-cytotoxic concentrations ranging from 35 to 100 µM (6, 7, 22, 23). The maximum non-cytotoxic concentration of NDGA in LLC-MK2 cells was determined to be 90.4 ± 5.9 µM (9). Antiviral activity against WNV was evaluated using a concentration range below this threshold to minimize host cell damage, resulting in a selectivity index (SI) of 3.9. The observed antiviral activity, combined with the absence of virucidal effects, is consistant with previous findings (7).

Time-of-addition experiments demonstrated that NDGA exerts its antiviral effect when added within the first 2 h PI. Considering that NDGA uptake into cells has already been demonstrated (9), these findings strongly suggest that NDGA demonstrates its antiviral activity intracellularly. Notably, this time-dependent antiviral effect was consistent with that observed against DENV1 (8), suggesting a shared mechanism of action against these related flaviviruses.

Flaviviruses, including WNV, are highly dependent on cellular lipid metabolism for replication. NDGA affects lipid metabolism primarily by inhibiting 5-LOX and SREBP pathways. The antiviral activity of NDGA has been previously linked to these inhibitory effects (6, 13). Merino-Ramos *et al*. (7) demonstrated that 35 µM of resveratrol, an inhibitor of SREBP1, exhibits antiviral activity against WNV, further supporting the role of SREBP inhibition in the antiviral mechanism of NDGA. Our findings suggest that inhibition of the 5-LOX pathway does not significantly affect WNV replication. In contrast, resveratrol (20 µM) modestly decreased residual WNV particles, although its antiviral effect was less pronounced compared to NDGA.

It is known that during the flavivirus replication cycle, specifically during the first hours of infection, the uncoating process of the viral particle occurs, which involves the release of the viral genome from the endosomes into the cytoplasm (24, 25). During this phase, viral particles are trafficked to late endosomes/lysosomes, where acidic pH triggers conformational changes in the viral envelope protein. This leads to membrane fusion and the release of the viral genome into the cytoplasm (25, 26). The trafficking process is critically dependent on the integrity of the cellular microtubule (MT) network and is tightly regulated by endosomal pH. MTs and their associated motor proteins play pivotal roles in the intracellular trafficking of viral particles within infected cells (27). Previous studies have investigated the role of MTs and the motor protein dynein in the early stages of DENV2 replication in BHK-21 cells (28). These studies demonstrated that the envelope (E) protein of DENV2 interacts with α-tubulin, suggesting that newly synthesized viral proteins are transported along MT tracks. Besides, dynein was found to be associated with the E protein of DENV2, facilitating its transport into the perinuclear region. Disruption of the dynein complex through dynamitin overexpression significantly impaired the trafficking of DENV E and core proteins (28). Furthermore, investigations on the trafficking of WNV E and C (capsid) proteins revealed their coordinated transport along MTs from the perinuclear region towards the plasma membrane. This process involves the motor protein kinesin and can be effectively inhibited by vinblastine sulfate, a well-known MT-disrupting agent (Chu and Ng, 2002). These results collectively underscore the critical role of the MT cellular network in supporting various stages of flavivirus replication, suggesting that disrupting MT function could represent a promising antiviral strategy. Cultures treated with NDGA for 2 and 24 h exhibited significant disruptions and alterations in the distribution of cellular MTs. Specifically, at 2 h post-treatment, MTs lost their typical radial organization from the microtubule-organizing center and adopted an irregular distribution toward the center of the cell. At 24 h, circular rearrangements of MTs became evident, suggesting a progressive and dynamic remodeling of the MT network over time. This observation is novel, as no previous reports have documented that NDGA induces such alterations in the MT fibers.

These observations align with existing research indicating that NDGA disrupts MT polymerization, leading to a reduction in MT density and significant alterations in MT organization (29). However, Nakamura et al. described NDGA as a weak microtubule-stabilizing agent that preserved the radial MTs organization and did not induce bundling, suggesting a more subtle effect on cytoskeletal architecture. In contrast, our findings demonstrate a marked reorganization of the MTs network, including loss of radial symmetry and atypical structural patterns not previously documented. These disruptions could impair cell motility and affect essential processes such as division and intracellular transport. Moreover, the influence of NDGA on tubulin dynamics may activate signaling pathways involved in apoptosis, underscoring its potential as a therapeutic agent in targeting cancer cell behavior. Overall, our results emphasize the significance of NDGA action on the tubulin cytoskeleton in modulating cellular functions and responses.

As already demonstrated by our group, NDGA possesses lysosomotropic effects, probably by disrupting the acidic environment necessary for viral envelope fusion within endosomes (8). The observed alterations in MTs could further influence viral replication by interfering with the transport of viral components, particularly during the crucial uncoating process.

Thus, the dual impact of NDGA on both endosomal acidification and cytoskeletal integrity may act simultaneously to impair early viral replication events.Collectively, these findings suggest that the antiviral activity of NDGA may result from a combination of effects on cellular machinery, including disruption of both endosome pH and the MT network, ultimately inhibiting viral replication.

In conclusion, our results reveal novel insights into the mechanism of NDGA antiviral activity, demonstrating its ability to disrupt critical cellular processes required for WNV replication. By targeting both the pH of the endosome and the MT network, NDGA offers a multifaceted approach to inhibit viral infection. These findings not only corroborate previous research but also significantly expand our understanding of the complex interactions between viruses and host cells. Taken together, these data provide a solid foundation for further research on the therapeutic potential of NDGA against Flavivirus infections.

## Material and Methods

### Cell Culture and Virus

LLC-MK2 (*Macaca mulatta* monkey kidney cells ATCC® CCL-7) were maintained in Eagle’s minimal essential medium (EMEM, Gibco, Grand Island, NY, USA) supplemented with 10% fetal bovine serum (FBS, Natocor, Carlos Paz, Argentina), 30 μg/mL of L-glutamine (Sigma-Aldrich) and 50 μg/mL gentamicin (Sigma-Aldrich, St. Louis, MO, USA). Cells were grown at 37 °C in a humidified atmosphere containing 5% CO_2_. The viral stock of WNV was the strain E/7229/062, which was titrated by the plaque-forming units (PFU) reduction method (30).

### Compounds and Standard Solutions

NDGA, with 95% purity, was extracted and isolated from *Larrea divaricata* Cav. (*Zigophyllaceae*) as we previously reported (8). Ribavirin (RIB, Filaxis, Buenos Aires, Argentina) was used as a positive antiviral control. Stock solutions of NDGA, resveratrol (Sigma-Aldrich), caffeic acid (Sigma-Aldrich), and RIB were prepared in dimethyl sulfoxide (DMSO, Tetrahedron, Mendoza, Argentina). Working solutions were subsequently prepared by diluting the stock solutions in EMEM supplemented with 2% of FBS, 30 μg/mL of L-glutamine (Sigma-Aldrich), and 50 μg/mL of gentamicin. The final DMSO concentration in all experiments was kept below 1%.

### Cell Viability Assay

The cytotoxic effects of NDGA on LLC-MK2 cells were assessed using the neutral red uptake and MTT (3-(4,5-dimethylthiazole-2-il)-2,5-diphenyltetrazolium) reduction assays as previously described (9). This allowed the calculation of the following parameters: CC_50_ (concentration causing 50% cell death), subtoxic concentration (SubTC, concentration affecting 20% of cells), and MNCC (maximum non-cytotoxic concentration).

### Synchronized Infection

A synchronized infection protocol was employed as previously described (8). Briefly, LLC-MK2 monolayers were infected with WNV at a multiplicity of infection (MOI) of 1. The virus adsorption was conducted by incubating at 4 °C for 1 h. Unbound virus was subsequently removed by washing with phosphate-buffered saline (PBS) at 4 °C. Cells were then incubated in EMEM with 2% FBS at 37 °C, 5% CO_2_, and 98% humidity. Cytopathic effect was examined through phase-contrast microscopy at low magnification with an Olympus IX81 microscope.

### Viral Replication Kinetic Curve

The cell monolayer was exposed to WNV at a MOI of 1. Then, synchronization of infection was performed to assess the length of the WNV replication cycle in LLC-MK2 cells. Subsequently, supernatants were collected at different times post-internalization (PI = 0 to 90 h), and viral particle titters were measured using the PFU assay (9).

### In Vitro Assessment of Virucidal Activity

To assess the direct virucidal activity of NDGA, the viral stock was incubated with NDGA in a 1:1 ratio (at a CC_50_ concentration of 115.7 µM) for 1 h at 37 °C. Control groups consisted of virus not exposed to NDGA (viral control, VC) and NDGA alone without virus (compound control). Ten-fold serial dilutions of each group were seeded on a confluent LLC-MK2 cell monolayer for 1 h at 37 °C to allow virus adsorption. A semisolid overlay was added, and the plates were incubated until plaque formation (5 days). Plaques were observed by staining with crystal violet (Anedra, Buenos Aires, Argentina) following fixation with 10% formalin (Anedra). Infectivity after treatment was determined by titration of residual virus using the PFU assay (31).

### In Vitro Antiviral Assay

A synchronized infection was performed in LLC-MK2 cells with a WNV suspension (100 PFU). Afterward, the infected cells were overlaid with EMEM and incubated at 37 °C for 1 h. After removing the inoculum, cells were treated with NDGA (2.5 to 90 µM) or with the positive control (+C), RIB (at non-cytotoxic concentration: 90 µM), all prepared in a semisolid medium and incubated for five days. Uninfected cells treated with NDGA were used as the cytotoxicity control (TC), whereas infected cells without treatment served as VC. Residual PFU in each condition was determined by plaque assay, and the inhibition percentage was calculated from the virus control (100% viable virus). The EC_50_ value, representing the concentration of NDGA required to inhibit 50% of viral replication, was determined from a dose-response curve. RIB was selected as +C for its application in treating other RNA virus infections (32).

### Time of Addition Assays

LLC-MK2 cells were pre-incubated with NDGA (90 μM) for 1 h at 37 °C (Pre-teatment, a). Subsequently, cells were washed twice with PBS and infected with WNV (100 PFU). The effect of NDGA at different stages of the viral replication cycle, including adsorption (AD, b), internalization (IN, c), and 1 h PI (d), was investigated following the methodology detailed by Martinez *et al*., (9). For conditions b and c, NDGA was added for 1 h and subsequently removed by washing with PBS. Semisolid medium was added in the three first conditions (a, b, and c), for final incubation at 37 °C with 5% CO_2_ until PFU formation. In condition d), NDGA was added at 1 h PI with the semisolid medium, followed by a final incubation until PFU formation. To further investigate the effect of the compound on the WNV replication cycle, NDGA (90 μM) was added simultaneously with the semisolid medium at different times PI (every hour for 8 h). The cells were then incubated at 37 °C with 5% CO_2_ for 5 days. Each experimental condition included VC, TC, and +C. Residual virus in all conditions were quantified by PFU, and treatments at different times were compared to VC.

### Effect of cellular lipid metabolism in WNV infection

To assess the antiviral efficacy of lipogenic pathway inhibitors against WNV, we employed the previously described methodology using non-toxic concentrations (9). The selected inhibitors were resveratrol targets sterol regulatory element binding protein 1 (SREBP1) (7) and caffeic acid inhibits 5-lipoxygenase (5-LOX) (18).

### Immunofluorescence assay

Initially, the immunofluorescence (IF) assay was set up using different concentrations of primary antibody from the serum of a patient in whom WNV had been confirmed. For this, cell monolayers were grown at 70% confluence on glass coverslips, infected with WNV at MOI 1 and treated with NDGA (90 μM) 1 h PI and the corresponding untreated control. After 12 and 24 h PI, monolayers were washed three times with PBS, fixed with 4% paraformaldehyde/1% sucrose (Riedel-de Haën, Sigma-Aldrich Laborchemikalien GmbH, Seelze, Germany) for 20 min at room temperature (RT). In parallel, to determine the effects of NDGA on the host cell cytoskeleton, cell monolayers were incubated for 2 and 24 h with NDGA (90 μM) and fixed with 4% paraformaldehyde/1% sucrose. Subsequently, the fixed cells were permeabilized with 0.2% TritonTM X-100 (Sigma-Aldrich) in PBS for 5 min at RT and incubated with 5% bovine serum albumin (BSA, Sigma-Aldrich) for 1 h at RT. To visualize WNV, the primary antibody was added at a 1/500 dilution in 1% bovine serum albumin overnight at 4 °C. The visualization procedure was performed using a mouse anti-α-tubulin primary antibodies (DM1A, Sigma-Aldrich) diluted 1/1000 in 1% BSA/PBS solution overnight at 4 °C. Subsequently, monolayers were washed three times with PBS and incubated with secondary antibodies (FITC-anti-human IgG or goat anti-mouse Alexa Fluor 488-# A-11017, Thermo Fisher Scientific Inc., Waltham, Massachusetts, USA) for 1 h at RT. Fixed and permeabilized cell monolayers were stained with Phalloidin-Tetramethyl rhodamine B (Sigma-Aldrich) for 1 h at RT to visualize the actin filaments (33).

Cells were observed using a conventional inverted epifluorescence microscope (Olympus IX81, Olympus Corporation, Shinjuku, Tokyo). Images were processed using Adobe Photoshop CS6 (Version: 13.0). The percentage of infected cells was calculated as the number of cells positive for the viral antigen from 10 microscope fields, selected at random, with comparable numbers of cells (more than 300 cells) per condition in each independent experiment. The percentage of infected cells in each treatment was determined by normalizing data compared to controls (100% value). To evaluate NDGA-induced morphological alterations in the tubulin cytoskeleton, the number of cells displaying structural abnormalities, characterized by cell rounding, disruption of the microtubule network, and perinuclear redistribution or condensation of tubulin fibers was quantified at 2 and 24 h post-incubation. For each condition, 300 cells were counted across randomly selected fields. The percentage of affected cells was normalized to the untreated control, which was set as 100%.

### Statistical Analysis

All statistical analyses and graphical representations were performed using GraphPad Prism 5. Data were expressed as mean ± standard deviation 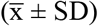, derived from three independent experiments, each with three inter-trial replicates. Statistical analyses were carried out using an unpaired Student T-test and one-way ANOVA test, followed by a Tukey post-hoc analysis to enable specific group comparisons (Type I error set at 0.05), as appropriate.

## Abbreviations

5-LOX: 5-lipoxygenase
+C: positive control
AD: adsorption
BSA: bovine serum albumin
CC_50_: cytotoxic concentration for 50% of the cells
DENV: Dengue virus
DMSO: dimethyl sulfoxide
EC_50_: effective concentration 50
EMEM: Eagle’s minimal essential medium
FBS: fetal bovine serum
IF: immunofluorescence
IN: internalization
MNCC: maximum non-cytotoxic concentration
MOI: multiplicity of infection
MTT: 3-(4,5-dimetiltiazol-2-il)-2,5-difeniltetrazolio
NDGA: nordihydroguaiaretic acid
PBS: phosphate-bufferd saline
PFU: plaque-forming units
PI: post-internalization
PT: pre-treatment
RIB: ribavirin
RT: room temperature
SI: selectivity index
SREBP: sterols regulatory elements binding proteins
SubTC: Subtoxic concentration
TC: toxicity control
VC: viral control
WNV: West Nile virus

## Acknowledgements

We appreciate the contribution of Dr. Carlos Tonn (INTEQUI-CONICET) for providing the pure nordihydroguaiaretic acid. This research was funded by: SECyT (Consolidar, tipo 2, s/res. N° 411/18 y 155/22), FONCyT (PICT 2018 N° 4576, s/ res. ANPCyT n° 401/19), CONICET (PIP 2021–2023, s/ res. 1639/2021) and SeCyT-UNC (Proyecto “FORMAR” s/Resol. 2020-233-E-UNC-SECYT#ACTIP).

## Authors’ contributions

**Florencia Martinez:** Writing original draft, Visualization, Validation, Data curation, Conceptualization, Investigation. **Lucia María Ghietto:** Writing original draft, Visualization, Formal analysis, Data curation, Conceptualization. **Juliana Lingua**: Writing original draft, Visualization. **Juan Javier Aguilar:** Writing original draft, Conceptualization. **Pedro Ignacio Gil:** Writing original draft, Data curation, Formal analysis. **Tomas Isaac Gomez:** Writing original draft, Visualization. **Juliana Marioni:** Writing original draft, Visualization. **Susana C. Núñez-Montoya:** Writing original draft, Conceptualization, Visualization, Funding acquisition. **Brenda S. Konigheim:** Writing original draft, Conceptualization, Visualization, Supervision, Funding acquisition.

## Data availability

Data will be made available on request.

## References

1. Bialosuknia SM, Dupuis Ii AP, Zink SD, Koetzner CA, Maffei JG, Owen JC, Landwerlen H, Kramer LD, Ciota AT. 2022. Adaptive evolution of West Nile virus facilitated increased transmissibility and prevalence in New York State. Emerg Microbes Infect 11:988–999.

2. Post LA, Lin JS, Moss CB, Murphy RL, Ison MG, Achenbach CJ, Resnick D, Singh LN, White J, Boctor MJ, Welch SB, Oehmke JF. 2021. SARS-CoV-2 Wave Two Surveillance in East Asia and the Pacific: Longitudinal Trend Analysis. J Med Internet Res 23:e25454.

3. Koch RT, Erazo D, Folly AJ, Johnson N, Dellicour S, Grubaugh ND, Vogels CBF. 2024. Genomic epidemiology of West Nile virus in Europe. One Health 18:100664.

4. Mingo-Casas P, Blazquez AB, Gomez de Cedron M, San-Felix A, Molina S, Escribano-Romero E, Calvo-Pinilla E, Jimenez de Oya N, Ramirez de Molina A, Saiz JC, Perez-Perez MJ, Martin-Acebes MA. 2023. Glycolytic shift during West Nile virus infection provides new therapeutic opportunities. J Neuroinflammation 20:217.

5. Manda G, Rojo AI, Martinez-Klimova E, Pedraza-Chaverri J, Cuadrado A. 2020. Nordihydroguaiaretic Acid: From Herbal Medicine to Clinical Development for Cancer and Chronic Diseases. Front Pharmacol 11:151.

6. Soto-Acosta R, Bautista-Carbajal P, Syed GH, Siddiqui A, Del Angel RM. 2014. Nordihydroguaiaretic acid (NDGA) inhibits replication and viral morphogenesis of dengue virus. Antiviral Res 109:132–40.

7. Merino-Ramos T, Jimenez de Oya N, Saiz JC, Martin-Acebes MA. 2017. Antiviral Activity of Nordihydroguaiaretic Acid and Its Derivative Tetra-O-Methyl Nordihydroguaiaretic Acid against West Nile Virus and Zika Virus. Antimicrob Agents Chemother 61.

8. Martinez F, Ghietto LM, Lingua G, Mugas ML, Aguilar JJ, Gil P, Pisano MB, Marioni J, Paglini MG, Contigiani MS, Nunez-Montoya SC, Konigheim BS. 2022. New insights into the antiviral activity of nordihydroguaiaretic acid: Inhibition of dengue virus serotype 1 replication. Phytomedicine 106:154424.

9. Martinez F, Mugas ML, Aguilar JJ, Marioni J, Contigiani MS, Nunez Montoya SC, Konigheim BS. 2021. First report of antiviral activity of nordihydroguaiaretic acid against Fort Sherman virus (Orthobunyavirus). Antiviral Res 187:104976.

10. Mahajan S, Choudhary S, Kumar P, Tomar S. 2021. Antiviral strategies targeting host factors and mechanisms obliging +ssRNA viral pathogens. Bioorg Med Chem 46:116356.

11. Gnabre JN, Brady JN, Clanton DJ, Ito Y, Dittmer J, Bates RB, Huang RC. 1995. Inhibition of human immunodeficiency virus type 1 transcription and replication by DNA sequence-selective plant lignans. Proc Natl Acad Sci U S A 92:11239–43.

12. Uchide N, Ohyama K, Bessho T, Toyoda H. 2005. Inhibition of influenza-virus-induced apoptosis in chorion cells of human fetal membranes by nordihydroguaiaretic Acid. Intervirology 48:336–40.

13. Syed GH, Siddiqui A. 2011. Effects of hypolipidemic agent nordihydroguaiaretic acid on lipid droplets and hepatitis C virus. Hepatology 54:1936–46.

14. Konigheim BS, Aguilar JJ, Grasso S, Contigiani MS, Núñez Montoya SC. 2012. Incidence of the nordihydroguaiaretic acid content on the in vitro antiviral activity of extracts obtained from Larrea divaricata Cav. (Zygophyllaceae). Latin American Journal of Pharmacy 31:659–664.

15. Mori M, Kovalenko L, Malancona S, Saladini F, De Forni D, Pires M, Humbert N, Real E, Botzanowski T, Cianferani S, Giannini A, Dasso Lang MC, Cugia G, Poddesu B, Lori F, Zazzi M, Harper S, Summa V, Mely Y, Botta M. 2018. Structure-Based Identification of HIV-1 Nucleocapsid Protein Inhibitors Active against Wild-Type and Drug-Resistant HIV-1 Strains. ACS Chem Biol 13:253–266.

16. Bhattacherjee P, Boughton-Smith NK, Follenfant RL, Garland LG, Higgs GA, Hodson HF, Islip PJ, Jackson WP, Moncada S, Payne AN, et al. 1988. The effects of a novel series of selective inhibitors of arachidonate 5-lipoxygenase on anaphylactic and inflammatory responses. Ann N Y Acad Sci 524:307–20.

17. Wang GL, Fu YC, Xu WC, Feng YQ, Fang SR, Zhou XH. 2009. Resveratrol inhibits the expression of SREBP1 in cell model of steatosis via Sirt1-FOXO1 signaling pathway. Biochem Biophys Res Commun 380:644–9.

18. Koshihara Y, Neichi T, Murota S, Lao A, Fujimoto Y, Tatsuno T. 1984. Caffeic acid is a selective inhibitor for leukotriene biosynthesis. Biochim Biophys Acta 792:92–7.

19. Kononchik JP, Jr., Hernandez R, Brown DT. 2011. An alternative pathway for alphavirus entry. Virol J 8:304.

20. Makino Y, Suzuki T, Hasebe R, Kimura T, Maeda A, Takahashi H, Sawa H. 2014. Establishment of tracking system for West Nile virus entry and evidence of microtubule involvement in particle transport. J Virol Methods 195:250–7.

21. Korsten C, Reemtsma H, Ziegler U, Fischer S, Tews BA, Groschup MH, Silaghi C, Vasic A, Holicki CM. 2023. Cellular co-infections of West Nile virus and Usutu virus influence virus growth kinetics. Virol J 20:234.

22. Koob TJ, Willis TA, Hernandez DJ. 2001. Biocompatibility of NDGA-polymerized collagen fibers. I. Evaluation of cytotoxicity with tendon fibroblasts in vitro. J Biomed Mater Res 56:31–9.

23. Villalobos-Sanchez E, Garcia-Ruiz D, Camacho-Villegas TA, Canales-Aguirre AA, Gutierrez-Ortega A, Munoz-Medina JE, Elizondo-Quiroga DE. 2023. In Vitro Antiviral Activity of Nordihydroguaiaretic Acid against SARS-CoV-2. Viruses 15.

24. Byk LA, Iglesias NG, De Maio FA, Gebhard LG, Rossi M, Gamarnik AV. 2016. Dengue Virus Genome Uncoating Requires Ubiquitination. mBio 7.

25. Barrows NJ, Anglero-Rodriguez Y, Kim B, Jamison SF, Le Sommer C, McGee CE, Pearson JL, Dimopoulos G, Ascano M, Bradrick SS, Garcia-Blanco MA. 2019. Dual roles for the ER membrane protein complex in flavivirus infection: viral entry and protein biogenesis. Sci Rep 9:9711.

26. White JM, Whittaker GR. 2016. Fusion of Enveloped Viruses in Endosomes. Traffic 17:593–614.

27. Greber UF, Way M. 2006. A superhighway to virus infection. Cell 124:741–54.

28. Shrivastava N, Sripada S, Kaur J, Shah PS, Cecilia D. 2011. Insights into the internalization and retrograde trafficking of Dengue 2 virus in BHK-21 cells. PLoS One 6:e25229.

29. Nakamura M, Nakazawa J, Usui T, Osada H, Kono Y, Takatsuki A. 2003. Nordihydroguaiaretic acid, of a new family of microtubule-stabilizing agents, shows effects differentiated from paclitaxel. Biosci Biotechnol Biochem 67:151–7.

30. Morales MA, Barrandeguy M, Fabbri C, Garcia JB, Vissani A, Trono K, Gutierrez G, Pigretti S, Menchaca H, Garrido N, Taylor N, Fernandez F, Levis S, Enria D. 2006. West Nile virus isolation from equines in Argentina, 2006. Emerg Infect Dis 12:1559–61.

31. Cheng HY, Lin LT, Huang HH, Yang CM, Lin CC. 2008. Yin Chen Hao Tang, a Chinese prescription, inhibits both herpes simplex virus type-1 and type-2 infections in vitro. Antiviral Res 77:14–9.

32. Takhampunya R, Ubol S, Houng HS, Cameron CE, Padmanabhan R. 2006. Inhibition of dengue virus replication by mycophenolic acid and ribavirin. J Gen Virol 87:1947–1952.

33. Ghietto LM, Gil PI, Olmos Quinteros P, Gomez E, Piris FM, Kunda P, Contigiani M, Paglini MG. 2022. Members of Venezuelan Equine Encephalitis complex entry into host cells by clathrin-mediated endocytosis in a pH-dependent manner. Sci Rep 12:14556.

