## SUPPLEMENTARY MATERIAL for "Mechanistic characterization of the antiviral effects of Nordihydroguaiaretic Acid against West Nile Virus"

Running Head: Early NDGA Antiviral Activity Against WNV

Florencia Martinez and Lucia M. Ghietto contributed equally to this work. Author order was determined on the basis of seniority

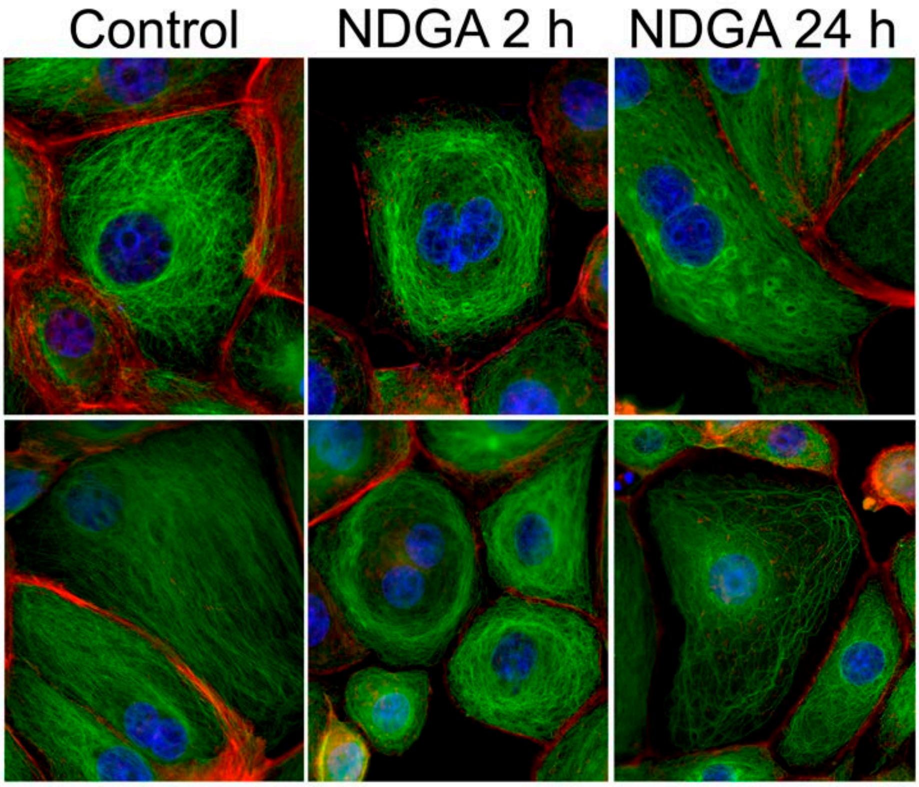


**Supplementary Figure S1. Morphological alterations of the tubulin cytoskeleton induced by NDGA.** Representative immunofluorescence images of Vero cells incubated with NDGA (90 µM) for 2 h and 24 h. Cells were fixed, permeabilized and stained with anti-α-tubulin antibody (green) and Phalloidin-Tetramethyl Rhodamine B to visualize actin filaments (red). NDGA treatment resulted in evident structural changes, including cell rounding, disruption of the microtubule network, and perinuclear condensation of tubulin filaments. These alterations were more pronounced at 24 h. Scale bar: 20 µm
